# Stable between-subject statistical inference from unstable within-subject functional connectivity estimates

**DOI:** 10.1101/268151

**Authors:** Diego Vidaurre, Mark W. Woolrich, Anderson M. Winkler, Theodoros Karapanagiotidis, Jonathan Smallwood, Thomas E. Nichols

**Affiliations:** Wellcome Trust Centre for Integrative Neuroimaging, Oxford Centre for Human Brain Activity, University of Oxford, UK.; Emotion and Development Branch, National Institute of Mental Health, National Institutes of Health, Bethesda, MD, USA.; Department of Psychiatry, Yale University School of Medicine, New Haven, CT, USA.; Department of Psychology, University of York, UK.; Big Data Institute, University of Oxford, UK

## Abstract

Spatial or temporal aspects of neural organisation are known to be important indices of how cognition is organised. However, measurements and estimations are often noisy and many of the algorithms used are probabilistic, which in combination have been argued to limit studies exploring the neural basis of specific aspects of cognition. Focusing on static and dynamic functional connectivity estimations, we propose to leverage this variability to improve statistical efficiency in relating these estimations to behaviour. To achieve this goal, we use a procedure based on permutation testing that provides a way of combining the results from many individual tests that refer to the same hypothesis. This is needed when testing a measure whose value is obtained from a noisy process, which can be repeated multiple times, referred to as replications. Focusing on functional connectivity, this noisy process can be: (i) computational, e.g. when using an approximate inference algorithm for which different runs can produce different results or (ii) observational, if we have the capacity to acquire data multiple times, and the different acquired data sets can be considered noisy examples of some underlying truth. In both cases, we are not interested in the individual replications but on the unobserved process generating each replication. In this note, we show how results can be combined instead of choosing just one of the estimated models. Using both simulations and real data, we show the benefits of this approach in practice.

## Introduction

Suppose that we are interested in testing hypotheses about variables, or set of variables, which we can observe on multiple occasions such that we may obtain a number of noisy measures of the same underlying (unobserved) feature or process. This can happen when we replicate a measurement on multiple occasions for each subject, or if the design of the experiment is such that the repetitions are independent from each other (which would not be the case, for example, if there is a strong effect of learning or habituation across runs). This can also happen when we are modelling data using an approach that is complex enough that inferences about the model parameters can be slightly different every time we estimate the model, e.g. with different arbitrary initialisations. This is the case, for example, for independent component analysis (ICA, Hyvärinen & Oja, 2000; Beckmann et al., 2005) and Hidden Markov models (HMM, Rabiner, 1989; Vidaurre et al., 2016).

In non-deterministic approaches such as ICA and HMM, the degree to which different initialisations will lead to different estimates (i.e. different local minima) of the model parameters depends on elements such as the signal-to-noise ratio, training parameters, and amount of available data (Himberg et al., 2004). Successive runs of the algorithm may find local minima that are equally good or equally likely, or it may find suboptimal local minima. While in some settings an appropriate figure of merit (e.g. residual sum of squares or model evidence) can adjudicate between these different estimates, sometimes no practical or definitive model comparison score is available; further, even when a score is available, this is typically an approximation or a heuristic, and it is possible that many models with very similar scores will be found. Here we claim that all models are potentially useful, and that an effective combination can be more powerful than choosing a single model. More specifically, in this work we take up the issue of making inference on these noisy replicate estimates, relating the estimates on a group of subjects to variables such as demographic, behaviour or personality scores. For this, we are not interested in whether each score relates to each individual replicate; rather, we aim to assess, based on a single global test over the pool of estimates, whether there is evidence that each score holds a significant association with the estimated measure.

Based on the principles of permutation testing, this paper presents a simple approach where we use the NPC algorithm (Pesarin and Salmaso, 2010; Winkler et al., 2016) to combine results from multiple functional connectivity (FC) estimations, regardless of whether the replications are at the level of data acquisition or model inference. This approach is useful in estimating effects that explain the underlying data that is the focus of the analysis. We demonstrate the validity of this method on the HMM, using simulations and data from the Human Connectome Project (Smith et al., 2013), where we test a measure of (resting state fMRI) dynamic FC over one hundred different HMM runs against a number of behavioural variables measured across hundreds of subjects.

## Methods

### Background

We refer to the noisy samples or parameter inference runs as R replications, to be distinguished from the P observed variables against which we aim to test. (Replications are not to be confused with realisations, which will use to refer to the multiple instances of the synthetic experimental scenario carried out below.) We have one hypothesis per observed variable, and wish to combine the tests across multiple replications, with no particular interest in assessing each replication in isolation. For N subjects, let us denote replications as Y (N by R), and observed variables as X (N by P). For reference, we will consider each column of Y (referred to as y_j_) as a noisy sample of certain unobservable variable of interest Y_0_.

For each column of Y and each column of X (referred to as x_i_), we can use permutations (Nichols and Holmes, 2002) to test the null hypothesis that there is no association between the model and the observed data. From this procedure we obtain a (1 by R) vector of p-values per observed variable, say p_j_. A simple approach could combine these R values with a simple statistic such as the mean or the median of p_j_ to assess the significance: if the mean p-value is small (e.g. below 0.01), this would suggest that there is a significant relationship between Y_0_ and x_j_. In what follows, we will refer to this summarised p-value as p_mean_, similar to Edgington’s p-value combining method comprised of the sum of p-values (Edington, 1972). A more effective approach is to use the geometric mean, equivalent to exponentiating the average of theWhick log p-values; this is related to Fisher’s p-value combining method (Fisher, 1932) and amplifies the importance of values near zero. Denoting the individual p-values for a given observed variable of interest as p_i_, we have

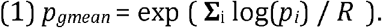

Again, if p_gmean_ is below a certain level, we can state there is a significant relationship between the replications and the examined observed variable. Note that neither p_mean_ or pgmean are p-values because they do not distribute uniformly in [0,1] under the null.

### Example case for a single pair of variables

Before coming to a complete description we consider a toy example to make the point above more concrete. We wish to assess if there is a linear relationship between two variables, a and b. The first one, a, with values a_n_, is Gaussian distributed (mean 0, standard deviation 1); the second one, b, is a corrupted version of a by the introduction of random noise:

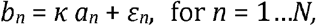

where ε_n_ are independent, Gaussian distributed random variables (mean 0, standard deviation 1), and κ ≥ 0. We generate replicates of b based on independent realisations of noise ε_n_ and κ, where κ is randomly sampled from a uniform distribution between 0 and c. We choose c to define the expected strength of the relationship between a and b. We then run permutation testing on each data set. We evaluate the power of the permutation combining method to detect a relationship between a and b for different values of c > 0. Even when κ is randomly small on some replicates, it may be large on others (allowing to detect the underlying relationship in these cases).

For the purpose of illustration we generated 10000 data sets using N=100, each with a different value of κ sampled from a uniform distribution, and performed permutation testing for each of them. We repeat this for three different values of c: 0.0, 0.1 and 0.2. Figure 1 shows histograms of correlation coefficients between a and b across data sets (top), and histograms of p-values (bottom). If the empirical distribution of p-values is basically flat, as is the case when c=0.0, then there no evidence of a relationship between a and b. However, when c=0.1 or c=0.2, then the distribution of p-values gets increasingly skewed toward zero. Therefore, if a and b were experimental replications of some pair of unobserved processes, we could intuitively say that there are signs of correlation between these processes in the c=0.1 and c=0.2 cases. However, neither p_mean_ or p_gmean_(not shown in the figure) are below 0.05; they are higher than 0.2 in all cases, emphasising again the point that p_mean_ or p_gmean_ are not p-values and, thus, the need for a permutation procedure to learn their null distribution.

**Figure 1.**
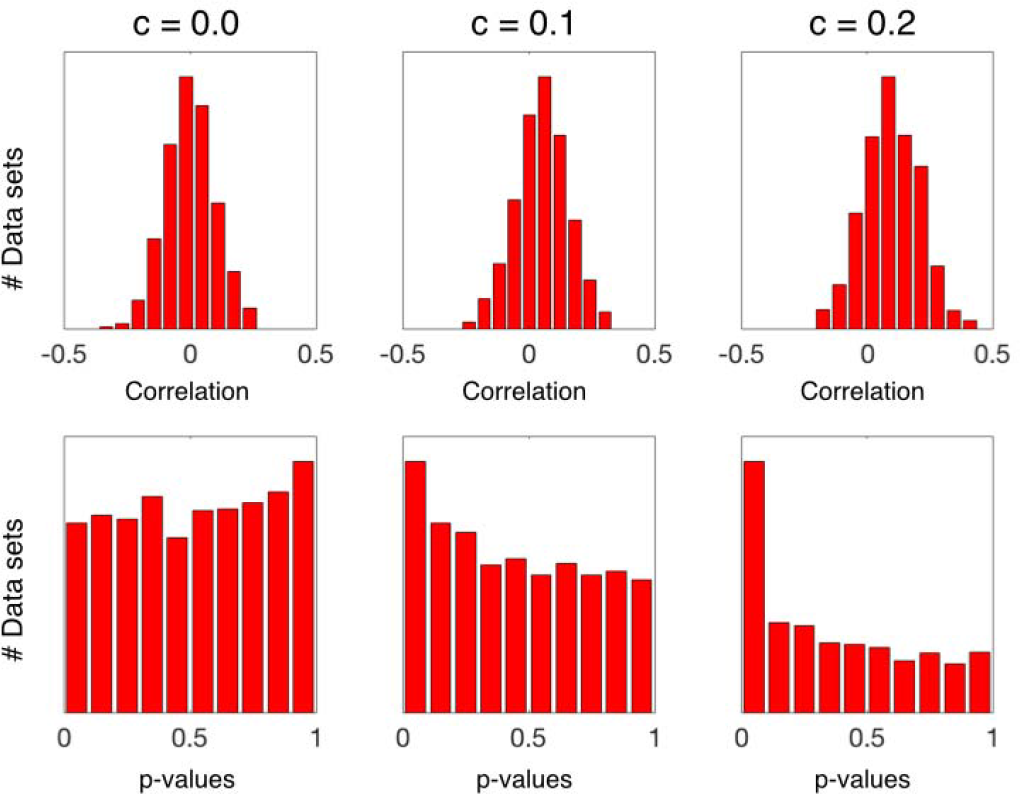
Distribution of correlation and (first-level) p-values for the toy example. Simulated examples where we generated 1000 data sets, where maximum regression coefficient, c, is systematically varied. When c>0.0, the mean correlation across data sets is higher than zero (top), and the distribution of p-values is skewed toward 0.0 (bottom). However, both pmean and pgmean are higher than 0.05.

### The NPC algorithm

We are interested on the relationship between the underlying variables (of which the replications are noisy observations), and not on the individual replications. Since p_mean_ or pgmean cannot be interpreted as p-values, we require a method to estimate actual p-values, that is, distributed uniformly under the null hypothesis. For this, we use the NPC algorithm on pgmean (Pesarin and Salmaso, 2010; Winkler et al., 2016). In the case when there is only one variable in the model (P=1), referred to as x, NPC (on p_gmean_) proceeds as follows:

I. Run statistical tests (e.g., t-tests) between each replication y_j_ and x to obtain an (R by 1) vector of p-values p^0^. We summarise p^0^ using the geometric mean, which, using Equation (1), yields pgmean. This corresponds to the first-level permutation testing.
II. Under the null hypothesis that each replication y_j_ and x are not associated, we randomly permute x a number of times K. For each permutation k, we produce an (R by 1) vector of parametric p-values p^k^ analogously to the previous step. We summarise p^k^ using the geometric mean, obtaining a surrogate 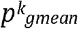 per permutation.
III. At the second level, we obtain a final p-value by computing the proportion of surrogate p-values 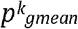 that are equal or lower than the unpermuted summary p-value pgmean:

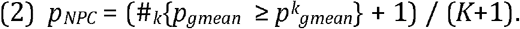

For the P > 1 case, that is, when there are more than one observed variable of interest, this procedure can be repeated for each variable, by using Equation (1) on the x_i_ separately. Crucially, we would use the same exact same permutations -that is. with the permutations happening in synchrony for all observed variables. This way, the dependence between the tests across variables is implicitly accounted for; in Winkler et al., 2016, this is referred to as “multiple models”. This will yield a final p-value per observed variable, say *p_NPC,J_*. We can obtain a summary, family-wise error corrected p-value (Nichols and Hayasaka, 2003) for each variable of interest j by computing

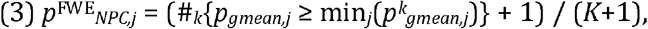

where 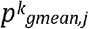,j is the null surrogate p-value obtained with Equation (1) for the j^th^ variable of interest and k^th^ realisation. Alternatively, we can use false-discovery rate (FDR; Benjamini and Hochberg, 1995; Nichols and Hayasaka, 2003) on the uncorrected p-values pNPC,j to obtain FDR-corrected p-values 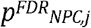.

In summary, this procedure draws statistical power from both working in logarithmic space (i.e. promoting the importance of p-values closer to zero), while simultaneously relaxing the alternative hypothesis from a highly conservative assumption – that “all of the replications bear a relationship with the corresponding observed variable” to a less conservative assumption that “at least some of the replications bear a relationship with the corresponding observed variable”. In the above example, for instance, this scheme of permutation testing produced a p-value higher than 0.5 when c=0.0, and p-values lower than 0.001 for both the c=0.1 and c=0.2 cases, exhibiting both sensitivity and robustness to non-normality (no distributional assumptions are made).

## Simulations

To illustrate the power of combining FC estimations using NPC, we simulated synthetic data sets emulating a scenario in which we are interested in testing whether FC between a pair of brain regions holds a relation to certain behavioural trait in a set of N subjects. In this situation, we have the following variables:

- A subject-specific FC coefficient β, which we cannot observe directly.
- A behavioural variable, hypothesised to be related to FC and encoded by a (N by 1) vector x, that can be observed directly.
- Some neural process modulated by β, denoted as S, which we cannot observe directly. We can consider S to be some archetypical, noiseless brain activity controlled by β.
- The observed (e.g. neuroimaging) data sets D, which are noisy measurements of S and have dimension (T by 2). This measurement can be repeated up to R times per subject.
- An (N by R) matrix Y, such that Y_nj_ contains the estimated FC value for the n^th^ subject and j^th^ experimental replication (i.e. the correlation coefficient between the channels of the corresponding measured data D).

In this context, the noise in the observations (or replications) stems from the imperfect measurement of S, which we can measure multiple times (R). Therefore, there is a relation between FC (β, which we cannot observe but we can estimate) and behaviour (x), but this relationship is noisy and weak for some replications. The objective of this simulation is then to assess whether the proposed approach can uncover such relationship. In detail, we generate data from this setting as follows:

We have N=200 subjects. We uniformly sample a value β_n_ between −0.2 and +0.2 for each subject n. For each subject, also, we sample two vectors with 10000 values each: the first, s_n1_, is Gaussian distributed (mean 0, standard deviation 1), whereas the second is set as

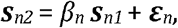

where ε_n_ is also Gaussian-distributed. The vectors s_n1_ and s_n2_ constitute the unobserved neural process S. The correlation between s_n1_ and s_n2_ can be analytically computed from β_n_ as

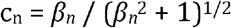

We set the value of the observed behavioural variable for each subject to be

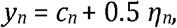

where η_n_ is Gaussian distributed (mean 0, standard deviation 1). Now, in order to sample the observed data sets D_n_ for each subject, we randomly sample 100 time points from S_n_ (whose columns are s_n1_ and s_n2_) and add some Gaussian noise with mean 0 and standard deviation σ. We do this R times per subject, obtaining one (100 by 2) noisy data set D_n_ =[d_n1_, d_n2_] each time. We then set the observed replication values to

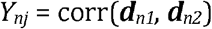

With both Y_n_ and x_n_ in hand, we run the described permutation testing algorithm on the noisily estimated FC matrix Y_n_ and the behavioural variable x_n_. By controlling σ (which defines how noisy are individual time series samples d_n1_ and d_n2_), we can make the detection more or less difficult. We use a range of 30 values for σ between 0.25 and 1.5, and repeat data generation and testing 100 times per value of σ. For each repetition of the experiment, standard permutation testing results on R=100 p-values (one per replication). Note that, since P=1, there is no need to control for familywise error rate across observed variables (Equation (3)).

On top, Figure 2 shows pmean / pgmean / pNPC (respectively from left to right) averaged across the 100 realisations of the experiment as a function of σ, together with 95% confidence intervals (minus/plus twice the standard error). Thanks to the effect of the logarithm, the pgmean values are lower than p_mean_ values, but neither of them ever reach significance provided the weak and volatile relationship between Y and x. The individual, per replication p-values (shown underneath for one example repetition, per value of σ) illustrate this point: although there are some significant p-values, the average is condemned to fail due to the frequent bad p-values associated to some too noisy replications. However, most of the p-values from the NPC permutation approach turned out to be significant despite the low magnitude of the signal across replications, with the average of p_NPC_ across realisations leaving the zone of significance only for the highest values of σ (i.e. for the hardest instantiations of the problem).

**Figure 2.**
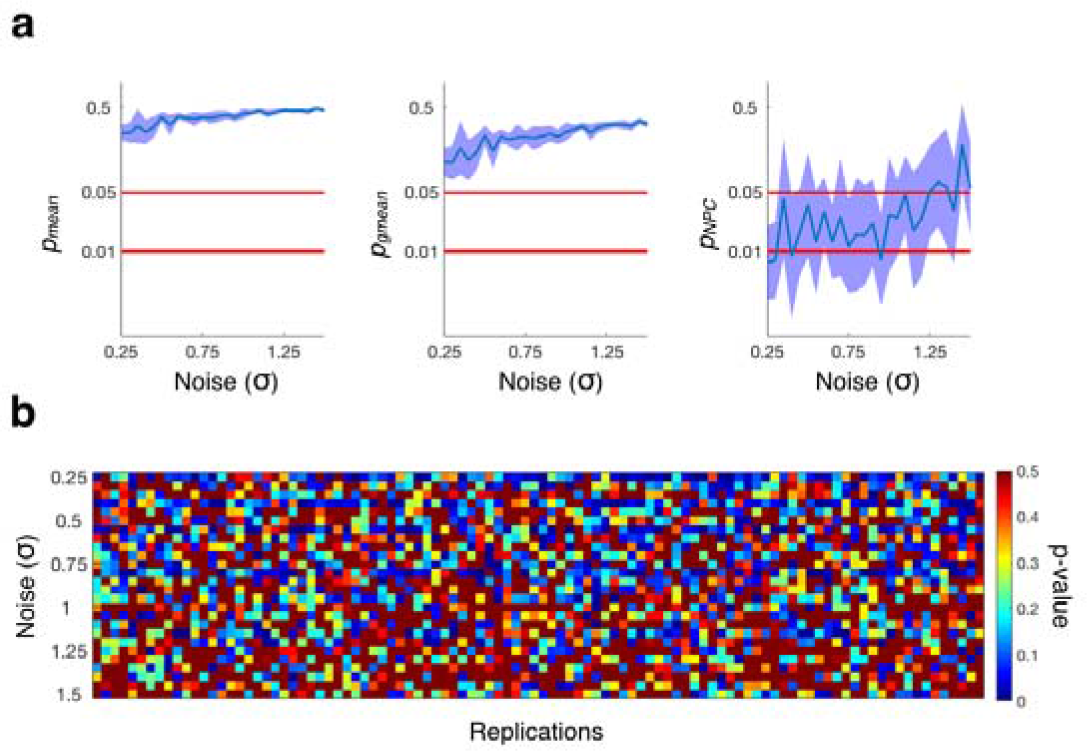
Results from the simulated data, where there is a relationship between the tested variables: FC and behaviour. (a) p-values obtained from combining tests by using the mean (pmean and pgmean), and p-values from the described permutation testing approach (p_NPC_), as a function of σ, which controls the noise in the replications (i.e. higher values of σ produce more difficult instantiations of the problem); intervals of confidence are computed across realisations of the experiment. (b) p-values before test combination for a given repetition (per value of σ).

Next, we repeat the same analysis but forcing a fixed value of β_n_ for all subjects (in particular, we set β_n_ = 0). In this case, there is relationship between behaviour and FC. Figure 3 shows that NPC, as well as the other methods, are robust and do not yield Type I errors in this scenario.

**Figure 3.**
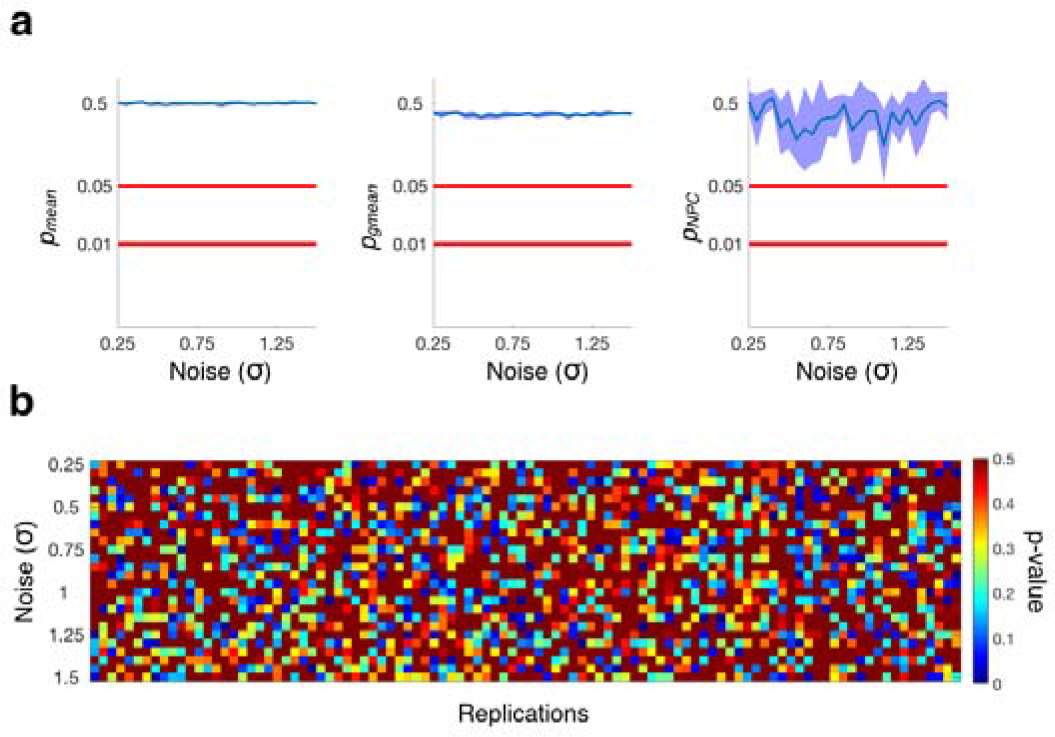
Results from the simulated data, where there is not a relationship between FC and behaviour. The description of the panels is equivalent to Figure 2. In this case, however, no relation was found between FC and behaviour, i.e. there are no Type I errors.

## Dynamic functional connectivity in real data

Having demonstrated the utility of the NPC approach to relate FC to behaviour in a synthetic scenario where the estimation was very noisy, we next evaluate it using real data by applying the Hidden Markov model (HMM) to resting state fMRI data from the Human Connectome Project (HCP). The HMM assumes that the data can be described using a finite number of states. Each state is represented using a probability distribution, which in this case is chosen to be a Gaussian distribution (Vidaurre et al, 2017a); that is, each state is described by a characteristic pattern of BOLD activation and a certain FC profile (we use the same configuration as in Vidaurre et al (2017a), to which we refer for further details). As the HMM is applied at the group level, the estimated states are shared across subjects; however, the state time courses that indicate the moments in time when each state is active are unique to a given individual. For the purposes of this analyses we set the HMM to have 12 states. Using the inferred state time courses, the amount of state-switching for each subject is calculated, which corresponds to a metric of how frequently subjects transition between different brain states (more specifically, given that the state time courses are probabilistic assignments, we compute the mean derivative of the state time courses for each subject). We use state switching as a summary metric of dynamic functional connectivity (DFC).

In order to infer the HMM at reasonable cost in spite of the large amount of data (820 subjects by 4 sessions by 15min, TR=0.75s), we use a stochastic learning procedure (Vidaurre et al, 2017b), which involves performing noisy, yet economical, updates during the inference. Since stochastic inference brings an additional layer of randomness into the HMM estimation but is not costly to run, we repeated the HMM inference 100 times and computed state-switching for each run. In this context, each HMM estimation constitutes a replication.

Furthermore, each subject has a number of behavioural measures, including psychological and sociological factors and several health-related markers. We used a total of 228 behavioural variables, after discarding those with more than 25% of missing values, in order to test against DFC as measure by state-switching. We included age, sex, motion and body-mass-index (the latter two usually considered as confounds). We also discarded those subjects without family information and those with a missing value in any of the behavioural variables. We used 10,000 permutations, respecting family structure of the HCP subjects (Winkler et al., 2015).

Although stochastic inference adds additional randomness to the estimation, the HMM has have previously been reported to perform very robustly in this data set (Vidaurre et al. 2017a), possibly as a consequence of the large number of subjects (N=820), the length of the scanning sessions, and the general high quality of the data. For this reason, the different HMM runs are quite consistent, which in turn means that the tests produce relatively similar results across replications (as shown below). To illustrate the effect of greater noise, we created a second set of replications where we permuted the state-switching measure between subjects randomly for half of the HMM runs (that is, half of the HMM runs, or replications, are potentially related to behaviour whereas the other half are noise, and all of them are included in the analysis). We refer to this as the perturbed data, as opposed to the original data where the HMM estimations are left intact.

Figure 4 compares the results of applying the NPC approach described above with the mean and geometric mean of the p-values, i.e. pmean and pgmean. Figure 4a shows the mean p-value (averaged across replications) reflecting the subject-wise correlation of state-switching (as measured by the HMM) with the different behavioural variables, with the behavioural variables being ordered from more to less significant; for purposes of illustration, dots represent individual p-values for some randomly chosen replications. On the left, the p-values obtained from standard permutation testing on the original HMM runs are quite consistent across replications; on the right, for the perturbed set of HMM runs, given that half were randomly ordered over subjects, the mean p-value reflects the reduced effect strength.

We next compute the cumulative sum of p-values for the 62 most significant variables (more specifically, those that had values less than 0.05 for all three methods, pmean, pgmean and NPC on pgmean). Figure 4b demonstrates the robustness of using NPC (on pgmean). On the left, where all the HMM runs were used normally, this difference is subtle; on the right, however, with perturbation the NPC’s p-values remain small while p_mean_ and pgmean grow large. The difference between p_mean_ and pgmean conveys the benefits of working on logarithm space, whereas the difference between pgmean and p_NPC_ reflects the transformation needed to convert pgmean to quantities interpretable as conventional p-values. Figure 4c shows, for each of the methods, the (combined across replications) p-values for the original data versus the perturbed data, reflecting that only the NPC approach is robust to having corrupted replications (i.e. the p-values are almost identical between the original and the perturbed data set).

**Figure 4.**
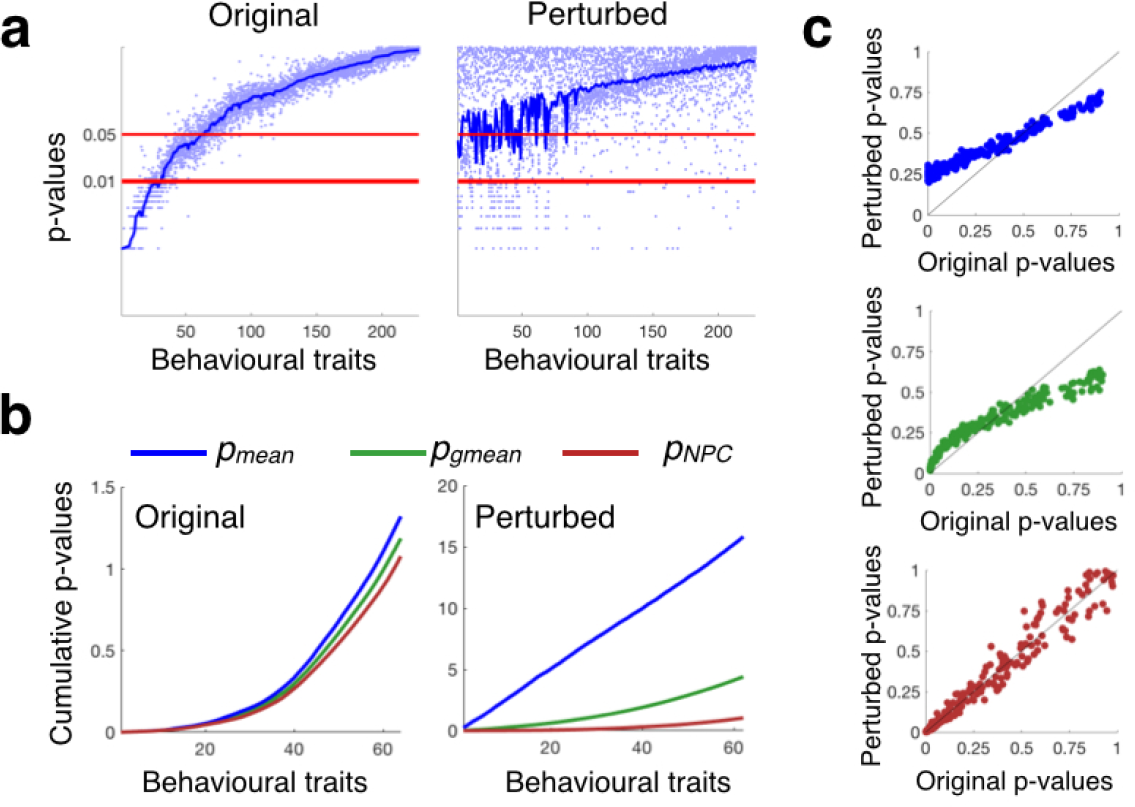
Analysis of the relation between behaviour and DFC (state switching) as measured by the HMM, where replications correspond to HMM runs. (a) Mean p-values (averaged over replications, with dots representing p-values for a randomly chosen subset of 10% of the individual replications), reflecting the subject-wise correlation of DFC with the different behavioural variables. On the X-axis, behavioural variables are ordered from more to less correlated. On the left, this is shown for the original data set; on the right, this is shown for the perturbed data set (a noisier version of the original data set). (b) NPC on pgmean outperforms p_mean_ and pgmean, as reflected when we examine the cumulative sum of p-values. (c) The p-values are robust to perturbation only for NPC, where correlation between perturbed and original p-values is close to 1.0.

Figure 5 presents the behavioural variables for which we found significance using the NPC procedure. Interestingly, although motion is a significant predictor it does not explain the greatest variance in this analysis, suggesting that DFC on resting state fMRI, as estimated by HMM, can be meaningfully related to behaviour beyond the influence of motion. Due to the relatively large number of observed variables, only a few were found to be significant after FWE correction (that is, in Equation (3), the minimum of the surrogate p-values across observed variables can be small if there are many observed variables to choose from). In contrast, FDR (Nichols & Hayasaka, 2003), allowed the identification of up to 26 variables. Importantly, if we randomly corrupt the entire data set (instead of half of the subjects as in the perturbed data set), all methods, including NPC, are able to avoid Type I errors by marking all behavioural variables as not significantly related to DFC (not shown).

**Figure 5.**
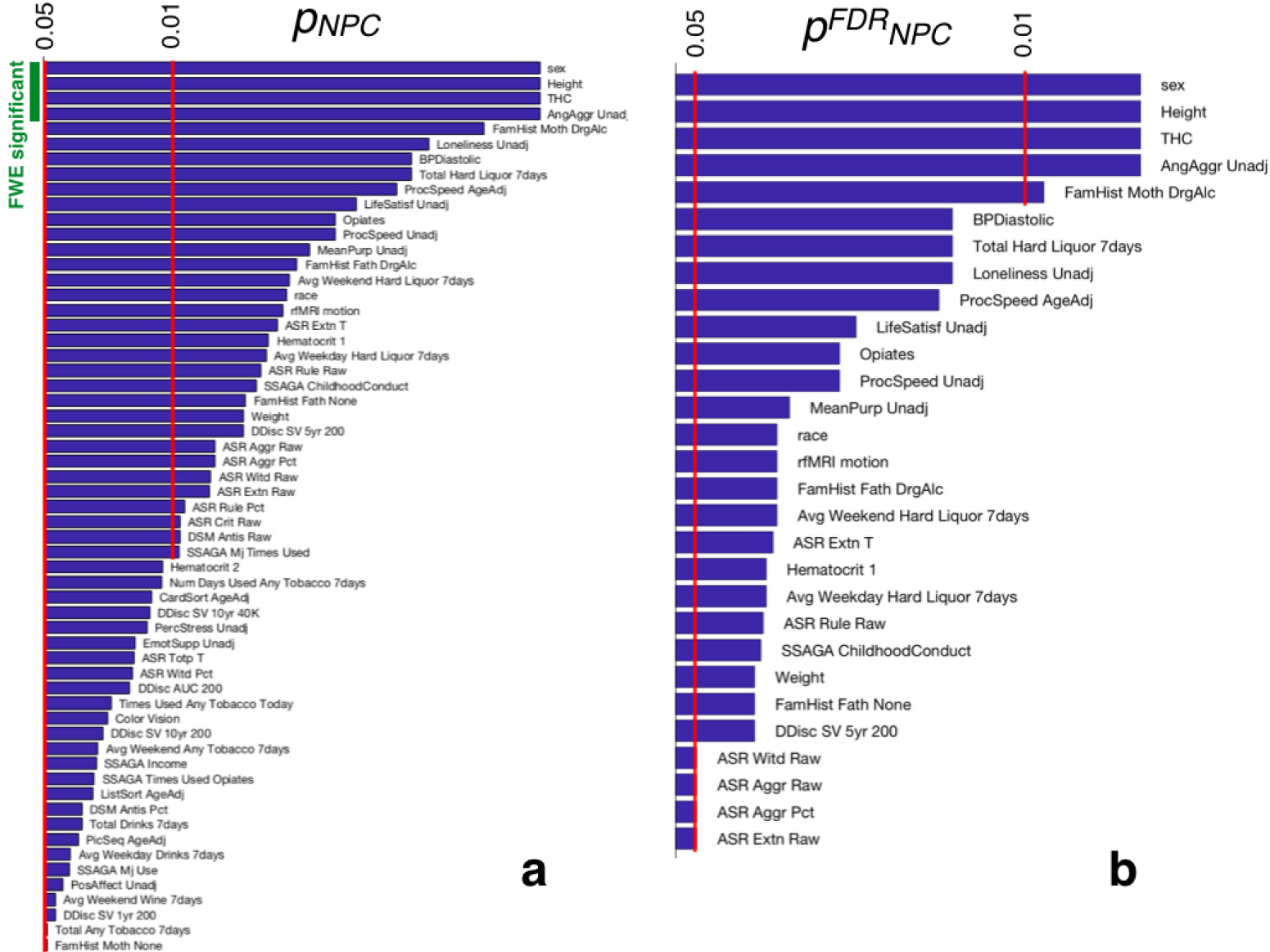
For the observed variables considered to be significant (out of 228), (a) p-values using the NPC on pgmean approach (p_NPC_), with FWE significance indicated on the top left; and (b) FDR-corrected p-values 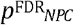.

## Discussion

In this paper, we show that the stochastic nature of FC estimations, often considered a hindrance, can be effectively integrated to provide valid and sensitive inferential proceedures. If the differences between the estimations are not only due to random noise, but contain different elements of information, such integration can be largely beneficial. If these differences are just pure noise, the presented procedure can approximate the accuracy of a single, noise-free estimation.

On these grounds, we describe a permutation testing approach based on previous work (Pesarin and Salmaso, 2010; Winkler et al., 2016) that can be used to test for the relationship between a set of observed variables and an unobserved (FC-based) variable for which we have a number of noisy estimations. The crucial point is that we are not interested in finding the relationships as described by a particular FC estimation, but instead would like to understand the relationship of the true FC with the observed variables. We took as a concrete example the relationship between covert patterns of intrinsic brain connectivity, as they occur at rest, and patterns of cognitive and demographic variables measured outside of the scanner, using data from the Human Connectome Project.

Although we focused on univariate observed variables and replications, the described method can straightforwardly be extended in a number of ways. First, although we focused on linear relationships between variables, it can easily be extended to multivariate statistics, such as multivariate linear regression, or canonical correlation analysis. This is important in that it allows studies in which the mapping between cognitive function and the data is not univariate in nature. It can also be extended to situations when we have replications on both sides of the correlation, such as when both the observed and non-observed behaviours are measured on multiple occasions. In this case, each pair of replications could be tested individually (for each element of the corresponding Cartesian product), and we would proceed similarly.

Moving forward, these types of approaches are likely to be particularly important in the domain of neuroscience given recent shifts towards the use of intrinsic connectivity at rest as a method of evaluating structural features of cognition. Intrinsic connectivity, as measured at rest, is a powerful tool for exploring the structure of neural organisation since it is able to reveal similar patterns of neural organisation as emerge during tasks (Smith et al., 2009). In addition, the simple non-invasive nature of the use of resting state as a method for assessing neural function means that it can be applied to multiple different populations, even those for whom task based measures of neural function, or psychological measurements may be problematic (such as children or populations with cognitive problems). Measuring neural organisation at rest is also easy to implement across centres making it amenable to the creation of large multicentre data sets, a shift that is likely to be increasingly important as neuroscience faces up to the challenges of reproducible science.

Despite the promise that assessing neural function at rest holds, many of the same features that make it an appealing tool for the cognitive neuroscience community are also at the heart of many of its limitations. For example, the power that is gained by the unobtrusive nature of the measure of neural function at rest also leads to concerns regarding what the measures actually represent: it is unclear which aspects of the neural signal reflect the intrinsic organisation of neural function, which reflect artefacts that emerge from physiological noise or motion (Power et al., 2013), and which reflect the patterns of ongoing experience that frequently emerge when individuals are not occupied by a demanding external task (Gorgolewski et al., 2014, Vatansever et al., 2017). In this context, because the underlying ground truth is unknown, an effective way to integrate estimations will help the researcher to identify which aspects of a given neural pattern are expressed in a robust way in relation to neurocognitive function.

Although dynamic approaches to understanding functional connectivity space are growing in popularity (Chang and Glover, 2010; Vidaurre et al., 2017a), different approaches have specific limitations. For example, sliding window approaches depend upon an apriori selection of the window length, which limits the granularity of neurocognitive states that can be identified. While approaches such as HMM circumvent this problem by allowing the data to determine the temporal duration of the underlying states, these analyses are inherently probabilistic and parameter inference can introduce noise into the analysis. In this context, NPC allows dynamic approaches to cognition to be compared to observed data in a systematic manner. This could help pave the way to formally evaluate how different descriptions of the underlying dynamics at rest best predict variables with well-described links to cognitive function. This way, NPC can become a useful tool in resolving the state-trait dichotomy that currently hinders the development of the science of how neural function evolves at rest.

## Acknowledgements

The Wellcome Centre for Integrative Neuroimaging is supported by core funding from the Wellcome Trust (203139/Z/16/Z). DV is supported by a Wellcome Trust Strategic Award (098369/Z/12/Z). MWW is supported by the Wellcome Trust (106183/Z/14/Z) and the MRC UK MEG Partnership Grant (MR/K005464/1). TEN is supported by the Wellcome Trust (100309/Z/12/Z).

## Bibliography

C.F. Beckmann, M. DeLuca, J.T. Devlin & S.M. Smith (2005). Investigations into resting-state connectivity using independent component analysis. Philosophical Transitions of the Royal Society B 360: 1001-1013.

Y. Benjamini & Y. Hochberg (1995). Controlling the false discovery rate: A practical and powerful approach for multiple testing. Journal of the Royal Statistical Society B, 57: 289-300.

C. Chang & G.H. Clover (2010). Time–frequency dynamics of resting-state brain connectivity measured with fMRI. NeuroImage 50, 81-98.

E.S. Edgington (1972): An additive method for combining probability values from independent experiments. The Journal of Psychology, 80, 351-363.

F.A. Fisher (1932). Statistical Methods for Research Workers, 4th ed. Edimburgh: Oliver Boyds.

K.J. Gorgolewski, D. Lurie, S. Urchs, J.A. Kipping, R.C. Craddock, M.P. Milham, D.S. Margulies & J. Smallwood (2014). A Correspondence between Individual Differences in the Brain’s Intrinsic Functional Architecture and the Content and Form of Self-Generated Thoughts. PLoS One 9, e97176.

A. Hyvarinen & E. Oja (2000). Independent component analysis: algorithms and applications. Neural Networks 13: 411-430.

J. Himberg, A. Hyvärinen & F. Exposito (2004). Validating the independent components of neuroimaging time series via clustering and visualization. NeuroImage 22, 1241-1222.

T.E. Nichols & S. Hayasaka (2003). Controlling the familywise error rate in functional neuroimaging: a comparative review. Statistical Methods in Medical Research, 12: 419–446.

T.E. Nichols & A.P. Holmes (2002). Nonparametric Permutation Tests For Functional Neuroimaging : A Primer with Examples, Human Brain Mapping, 15: 1–25.

F. Pesarin and L. Salmaso (2010). Permutation Tests for Complex Data: Theory, Applications and Software. West Sussex, England, UK: John Wiley and Sons.

L.R. Rabiner (1989). A tutorial on hidden Markov models and selected applications in speech recognition. Proceedings of the IEEE 77: 257-286.

S.M. Smith, C.F. Beckmann, J. Andersson, E.J. Auerbach, J. Bijsterbosch, G, Douaud, E. Duff, D.A. Feinberg, L. Griffanti, M.P. Harms, M. Kelly, T. Laumann, K.L. Miller, S. Moeller, S. Petersen, J. Power, G. Salimi-Khorshidi, A.Z. Snyder, A.T. Vu, M.W. Woolrich, J. Xu, E. Yacoub, K. Ugurbil, D.C. Van Essen & M.F. Glasser. (2013) Resting-state fMRI in the Human Connectome Project. NeuroImage 80: 144-168.

S.M. Smith, P.T. Fox, K.L. Miller, D.C. Glahn, P.M. Fox, C.E. Mackay, N. Filippini, K.E. Watkins, R. Toro, A.R. Laird & C.F. Beckmann (2009). Correspondence of the brain’s functional architecture during activation and rest. Proceedings of the National Academy of Sciences of the USA 106, 13040-13045.

D. Vatansever, D. Bzdok, H. Wang, G. Mollo, M. Sormaz, C. Murphy, T. Karapanagiotidis, J. Smallwood & E. Jefferies (2017). Varieties of semantic cognition revealed through simultaneous decomposition of intrinsic brain connectivity and behaviour. NeuroImage 158, 1-11.

D. Vidaurre, A.J. Quinn, A.P. Baker, D. Dupret, A. Tejero-Cantero & M.W. Woolrich (2016). Spectrally resolved fast transient brain states in electrophysiological data. NeuroImage 126: 81-95.

D. Vidaurre, R. Abeysuriya, R. Becker, A.J. Quinn, F. Alfaro-Almagro, S.M. Smith & M.W. Woolrich (2017b). Discovering dynamic brain networks from Big Data in rest and task. NeuroImage. In press.

D. Vidaurre, S.M. Smith & M.W. Woolrich (2017a). Brain networks are hierarchically organised in time. Proceedings of the National Academy of Sciences of the USA, In press.

M.J. Wainwright & M.I. Jordan (2008). Graphical models, exponential families and variational inference. Foundations and Trends in Machine Learning 1, 1-305.

A. Winkler, M.A. Webster, D. Vidaurre, T.E. Nichols & S.M. Smith (2015). Multi-level block permutation. NeuroImage 123: 253-268.

A. Winkler, M.A. Webster, J.C. Brooks, I. Tracey, S.M. Smith & T.E. Nichols (2016). Non-parametric combination and related permutation tests for Neuroimaging. Human Brain Mapping 37, 1486-1511.

